# The metabolic redox regime of *Pseudomonas putida* tunes its evolvability towards novel xenobiotic substrates

**DOI:** 10.1101/314989

**Authors:** Özlem Akkaya, Danilo R. Pérez-Pantoja, Belén Calles, Pablo I. Nikel, Victor de Lorenzo

## Abstract

During evolution of biodegradation pathways for xenobiotic compounds, the transition towards novel substrates of Rieske non-heme iron oxygenases borne by environmental bacteria is frequently associated with faulty reactions. Such reactions release reactive oxygen species (ROS), endowed with high mutagenic potential. The present work studies how the operation of a given metabolic network by a bacterial host may either foster or curtail the still-evolving biochemical pathway for catabolism of 2,4-dinitrotoluene (2,4-DNT). To this end, the genetically tractable strain *Pseudomonas putida* EM173 was chromosomally implanted with a Tn7 construct carrying the whole genetic complement (recruited from the environmental isolate *Burkholderia* sp. R34) necessary for complete biodegradation of 2,4-DNT. By using reporter technology and direct measurements of ROS formation, we observed that the engineered *P. putida* strain experienced oxidative stress when catabolizing the nitroaromatic substrate. However, ROS was neither translated into significant activation of the SOS response to DNA damage nor resulted in a mutagenic regime (unlike *Burkholderia* sp. R34, the original host of the pathway). To inspect whether the tolerance of *P. putida* to oxidative insults could be traced to its characteristic reductive redox regime, we artificially lowered the pool of NAD(P)H by conditional expression of a water forming, NADH-specific oxidase. Under the resulting low-NAD(P)H status, 2,4-DNT triggered a conspicuous mutagenic and genomic diversification scenario. These results indicate that the background biochemical network of environmental bacteria ultimately determines the evolvability of metabolic pathways. Moreover, the data explains the efficacy of some bacteria such as Pseudomonads to host and evolve new catabolic routes.

**IMPORTANCE:** Some environmental bacteria evolve new capacities for aerobic biodegradation of chemical pollutants by adapting pre-existing redox reactions to recently faced compounds. The process typically starts by co-option of enzymes of an available route to act on the chemical structure of the substrates-to-be. The critical bottleneck is generally the first biochemical step and most of the selective pressure operates on reshaping the initial reaction. In Rieske non-heme iron oxygenases, the interim uncoupling of the novel substrate to the old enzymes results in production of highly mutagenic ROS. In this work, we demonstrate that the background metabolic regime of the bacterium that hosts an evolving catabolic pathway (e.g. biodegradation of the xenobiotic 2,4-DNT) determines whether the cells would either adopt a genetic diversification regime or a robust ROS-tolerant state. These results expose new perspectives to contemporary attempts for rational assembly of whole-cell biocatalysts, as pursued by present-day metabolic engineering.

## INTRODUCTION

Environmental bacteria that catabolize xenobiotic pollutants existing only since the onset of synthetic chemistry offer a unique opportunity to inspect the rules that govern evolution of metabolic networks (1, 2). Unlike resistance to antibiotics, which can be caused by mutations modifying the target or by evolving just one protein (3-6), new catabolic phenotypes require multiple changes in the protein complement of the pathway along with other functions in the host that allow tuning of the novel route to the background biochemical network (7-10). Among microorganisms known to host aerobic routes for catabolism of typical industrial pollutants, e.g. chloroaromatic (11, 12) and nitroaromatic chemicals (13, 14), Pseudomonas species stand out as recurrent hosts of catabolic routes that enable growth on such unusual chemicals (15-20). This state of affairs raises two related questions. The first question is why new pathways evolve preferentially in this bacterial domain and not so much in other species. The second question is how an evolutionary solution to the metabolic challenge remains in the same bacterial domain rather than propagating into other prokaryotic hosts (15). The biochemical reactions at stake often involve redox transactions such as mono- or di-oxygenations executed by Rieske non-heme iron enzymes (21-23). Owing to their mechanism of action, when such oxygenases act on substrates-to-be that do not fit well in the active enzyme center, reactive oxygen species (ROS) are released (24-27). This phenomenon is due to a suboptimal kinetic scenario in which the substrate leaves the active pocket unscathed before the Fe-activated oxygen can attack the aromatic structure (28, 29). Uncoupled reactions of this sort result in considerable redox stress in the host (30) and—at least in some cases—DNA damage and acquisition of a mutagenic regime (31). The interplay between faulty redox reactions, ROS formation, and DNA damage has been studied in strain *Burkholderia* sp. R34. This environmental bacterium degrades (if with difficulties) 2,4-dinitrotoluene (2,4-DNT), an archetypal xenobiotic compound (32). The first enzyme of the dnt pathway is a Rieske non-heme iron dioxygenase that has evolved from a precursor protein acting on naphthalene (33). The extant catabolic pathway is still evolving, as its substrate profile and regulation keep features of the antecedent route. When *Burkholderia* sp. R34 is exposed to 2,4-DNT, the substrate is indeed degraded, but cells undergo a massive intracellular production of ROS (34) stemming from the first reaction (i.e. dioxygenation of the nitroaromatic compound to yield 4-methyl-5-nitrocatechol). While ROS kill most of the population, this DNA-damaging and protein-perturbing agent makes the surviving cells to diversify genetically (34). At least part of the ROS-triggered DNA mutagenesis can be traced to misincorporation of 8-hydroxy-2’-deoxyguanosine (8-oxoG) to DNA, although other mechanisms of ROS-mediated inhibition of the DNA mismatch repair system could be at play (35-37). The example of 2,4-DNT degradation illustrates how stress arising from an abortive metabolic reaction can paradoxically promote evolution of novel routes, as genetic diversification fosters exploration of the solution space by the whole bacterial population, plausibly leading to an optimized catabolic outcome. One consequence is that the evolution of aerobic degradation pathways for xenoaromatic compounds can occur only in bacterial hosts able to cope with intracellular ROS generation to a level that allows genetic diversification without trespassing a deadly threshold. The most common mechanism to counter oxidative stress involves the action of detoxifying enzymes (e.g. catalases, peroxidases, and hydroperoxide reductases) that inactivate ROS (38). The corresponding reactions are ultimately fed by metabolic NADPH (39, 40), which provides the reductive currency to counteract the noxious effects of ROS, e.g. via reduced glutathione (41, 42).

In this work, we addressed the effect of the background redox metabolism on evolvability of bacteria that host new biodegradation pathways. We have implanted the dnt route for catabolism of 2,4-DNT in the genome of the model soil bacterium *Pseudomonas putida*, and inspected the effect of metabolizing this compound on intracellular ROS production, redox stress, and genetic variability. *P. putida* is a frequent host of pathways for aerobic degradation of aromatics, and it is a habitual carrier of both evolving routes and naturally optimized pathways (43). These qualities are generally attributed to the distinct core metabolic network of this bacterium, geared to produce high NADPH levels (44, 45), further reinforced through the action of stress-induced pyridine nucleotide transhydrogenases (46). The results below indicate that ROS, resulting from faulty reactions of the dnt pathway on 2,4-DNT, are translated into genetic diversification of the host in a fashion that depends on its redox status—and therefore that evolvability of new traits is ultimately tuned by metabolism. Consequently, some bacteria seem to be more suitable for hosting evolution of new pathways and delivering their activities in sustained form.

## RESULTS

### Construction of a stable 2,4-DNT–degrading *P. putida* strain

The 2,4-DNT degradation pathway of *Burkholderia* sp. R34 converts the aromatic substrate into propionyl-coenzyme A and pyruvate through the sequential action of six enzymes (DntA to DntE, **Fig. 1a**). The eight genes encoding the entire pathway (i.e. dntAaAbAcAd dntB dntD dntE dntG), along with an ORF encoding a native regulatory protein (i.e. dntR) (47), were cloned from *Burkholderia* sp. R34, assembled in a synthetic Tn7-based vector (**Fig. 1b**), and delivered into the chromosome of *P. putida* EM173. This strain is a derivative of wild-type KT2440 devoid of four prophages and the endogenous Tn7 transposase (48), and displays enhanced genetic stability, a trait exploited when manipulating enzymes involved in harsh biochemical reactions (49). After checking the proper insertion of the genes into the target chromosome by PCR, the capability of the resulting strain (termed *P. putida* EM·DNT) to degrade 2,4-DNT was tested. For this, cells were grown overnight at 30°C and then diluted in fresh M9 minimal medium with 0.4% (wt/vol) succinate. When cultures reached an optical density at 600 nm (OD_600_) of 0.5, 2,4-DNT was added at 0.5 mM. After 3 h of incubation—and similarly to the original 2,4-DNT–degrading *Burkholderia* strain (34)—*P. putida* EM·DNT secreted distinctively colored metabolites (**Fig. 1c**), which indicated the activity of the dnt route in the surrogate *P. putida* host.

**Fig. 1.**
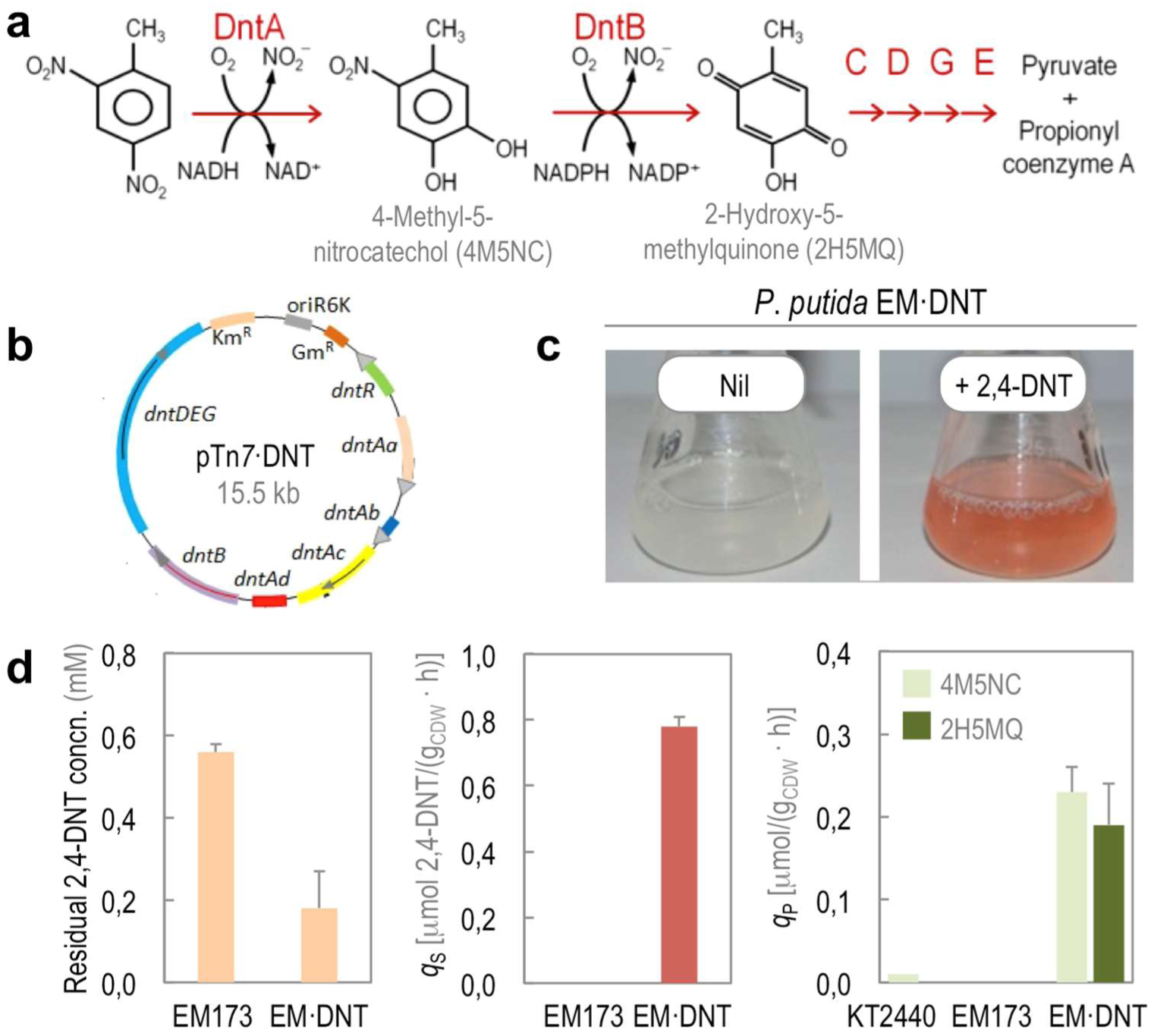
Construction and testing of a 2,4-DNT–degrading *P. putida* strain. **(a)** 2,4-DNT mineralization pathway from *Burkholderia* sp. R34. The catabolic route starts with the action of DntA, a 2,4-DNT dioxygenase belonging to the Rieske non-heme iron family that hydroxylates the aromatic ring in positions 4 and 5 to yield 4M5NC, releasing the first nitro substituent. The substituted catechol is subsequently mono-oxygenated by DntB, a 4M5NC hydroxylase that eliminates the remaining nitro group in the structure producing 2H5MQ. The rest of the pathway (executed by DntCDGE) includes a ring cleavage reaction and channeling of the products towards central carbon metabolism. **(b)** Assembly of the genes encoding the whole route for 2,4-DNT degradation, along with the *dntR*-encoded regulatory protein, in a synthetic Tn7 transposon. The resulting plasmid, pTn7·DNT, was delivered at the defined *att*Tn7 chromosomal site of the target host for stable insertion of the dnt gene cluster. **(c)** Qualitative testing of the recombinant *P. putida* EM·DNT strain, in which the dnt genes have been stably inserted in the chromosome of strain EM173. The appearance of a reddish color in cultures added with 2,4-DNT at 0.5 mM indicates the presence of 2H5MQ. **(d)** Quantification of kinetic parameters in cultures of *P. putida* KT2440 (wild-type strain), EM173 (a reduced-genome derivative of strain KT2440), and EM·DNT (expressing the *dnt* genes) grown in the presence of 2,4-DNT at 0.5 mM. The specific rates of 2,4-DNT consumption (q_*S*_) and formation of 4M5NC and 2H5MQ (q_*P*_) were calculated by measuring the concentration of the substrate and the products in culture supernatants. Bars represent mean values ± SD (*n* = 4) obtained after 24 h of incubation. Concn., concentration; CDW, cell dry weight.

As a quantitative measure of the activities implanted in *P. putida*, both the consumption of 2,4-DNT and the appearance of key metabolic intermediates in the route were assessed by gas chromatography coupled to mass spectrometry (**Fig. 1d**). After 24 h of incubation, *P. putida* EM·DNT processed 64% of the aromatic substrate, with a specific consumption rate of 0.78 ± 0.04 μmol 2,4-DNT · g cell dry weight (CDW)-1 · h-1. Expectedly, no substrate consumption was detected in cultures of the parental strain EM173. Both 4-methyl-5-nitrocatechol and 2-hydroxy-5-methylquinone, the metabolic products of DntA and DntB, respectively (50, 51), were detected in supernatants of *P. putida* EM·DNT cultures added with 2,4-DNT, but not in control experiments. The specific formation rates of 4-methyl-5-nitrocatechol and 2-hydroxy-5-methylquinone in these cultures were 0.24 ± 0.03 and 0.19 ± 0.05 μmol · g_CDW_^-1^ · h^-1^, respectively. Taken together, these results indicate that the 2,4-DNT degradation pathway grafted into *P. putida* was active under the culture conditions tested.

### 2,4-DNT degradation by *P. putida* EM·DNT results in ROS generation and activation of the cellular response to oxidative stress

Despite having the necessary genetic and biochemical complement, *Burkholderia* sp. R34 grows poorly on 2,4-DNT as the sole carbon source. This difficulty that can be traced to the formation of ROS upon exposure of the cells to the substrate of the biodegradation pathway (34). Against this background, we evaluated the generation of ROS in *P. putida* EM·DNT when exposed to 2,4-DNT at either 0.25 or 0.5 mM, assessing the fluorescence brought about by the ROS-sensitive dye 2’,7’-dichlorodihydrofluorescein diacetate (H2DCF-DA) in single cells by flow cytometry (**Fig. 2a**). The amount of ROS proportionally increased with respect to the concentration of 2,4-DNT, indicating that substrate consumption generates oxidative stress. The proportionality between the activity of the degradation pathway and ROS generation was qualitatively assessed in these experiments, since the amount of 2-hydroxy-5-methylquinone produced by the cells increased with the concentration of 2,4-DNT in the same fashion as ROS formation did. Quantification of ROS indicated that the mere exposure of the cells to 2,4-DNT is not the main cause of ROS generation; instead, the biochemical transformation of the substrate leads to endogenous oxidative stress. Indeed, *P. putida* EM·DNT had a ca. 10-fold increase in ROS formation upon exposure to 2,4-DNT (**Fig. 2b**). In contrast, strain EM173 had a lower accumulation of ROS (less than 4-fold increase) than strain EM·DNT when challenged with 2,4-DNT, even at the highest substrate concentration tested.

**Fig. 2.**
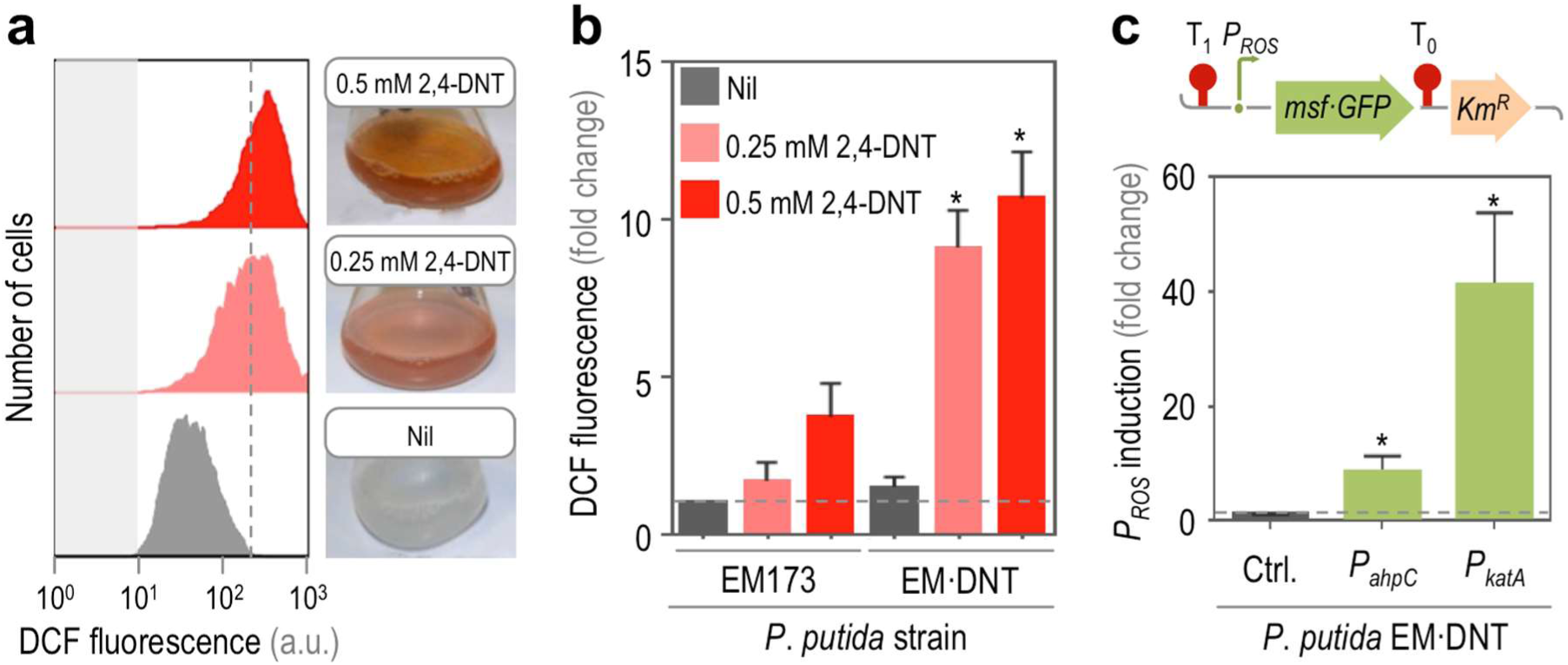
Phenotypic and transcriptional stress response of *P. putida* EM·DNT exposed to 2,4-DNT. **(a)** and **(b)**, Flow cytometry-assisted determination of ROS formation. **(a)** Histograms of raw data from untreated cells (Nil, added with DMSO, the 2,4-DNT solvent carrier), and cells exposed to 2,4-DNT at 0.25 or 0.5 mM. Cell suspensions were treated with the ROS-sensitive probe H_2_DCF-DA, and the resulting dichlorofluorescein (DCF) fluorescence levels were recorded in at least 15,000 individual cells. Gray rectangles indicate the maximum fluorescence in cells without addition of H_2_DCF-DA. A representative experiment per condition is shown in the diagram. **(b)** Fold change of the geometric mean (x-mean) of DCF fluorescence levels for each experimental condition in *P. putida* EM173 (parental strain) and EM·DNT (carries the dnt gene cluster). Bars represent the average of x-mean ± SD (*n* = 6), and the asterisk identifies significant differences at *P* < 0.05 as indicated by the Mann-Whitney U test. The dashed gray line indicates the basal level of DCF fluorescence in the control experiment. (c) Transcriptional activity of stress-responsive promoters. Two different oxidative stress reporters were constructed by placing the corresponding promoter (P_*ROS*_) in a pBBR1-based, kanamycin-resistant (Km^R^) vector bearing the promoterless gene encoding the monomeric and superfolder GFP (msf·GFP). The P_*ROS*_→msf·GFP construct was transcriptionally insulated by means of the T0 and T1 terminators. Elements not drawn to scale. The x-mean of the msf·GFP fluorescence was detected by flow cytometry in *P. putida* EM·DNT exposed to 2,4-DNT at 0.5 mM. The resulting msf·GFP fluorescence was compared to that in cells harvested from cultures that were not treated with 2,4-DNT (Ctrl., baseline indicated with a dashed gray line). Bars represent the mean value of the x-mean of the msf·GFP fluorescence ± SD (n = 4), and the asterisk identifies significant differences at *P* < 0.05 as indicated by Student’s *t* test.

Yet, is ROS formation connected to the activation of stress responses in the engineered *P. putida* strain? To answer this question, the transcriptional activity of genes involved in the oxidative stress response (i.e. ahpC, *PP_2439*, alkyl hydroperoxide reductase, and katA, *PP_0481*, a catalase) was studied by fusing the corresponding promoter region of these two genes to the reporter monomeric superfolder green fluorescent protein (msf·GFP, **Fig. 2c**). Either the ROS-reporter plasmids, or pSEVA237M, the promoter-less, msf·GFP-containing vector (52), were transformed into *P. putida* EM·DNT, and the msf·GFP signal was evaluated in cultures exposed to 2,4-DNT by flow cytometry. The output signal qualitatively followed the same trend as observed for ROS accumulation (**Fig. 2b**): the induction of the two oxidative stress-responsive promoters significantly increased in the presence of the aromatic compound (i.e. 10- and 42-fold increase in the transcriptional activity of P_*ahpC*_ and P_*katA*_, respectively, when cells were exposed to 2,4-DNT at 0.25 mM). In accordance with previous observations in *Burkholderia* sp. R34 (34), these results indicate that cells carrying the enzymes needed to process 2,4-DNT undergo oxidative stress conditions upon exposure to the substrate of the degradation pathway.

### Assessment of 2,4-DNT degradation on the SOS response and recombinogenic activity

Since the hypothesis underlying this work is that endogenous oxidative stress brought about by the biodegradation of an aromatic substrate could result in genetic novelty, we explored two types of mutagenic effects on the genome of *P. putida*. In the first place, and in order to evaluate if the degradation of 2,4-DNT promotes *recA*-mediated homologous DNA recombination, a reporter strain was designed as follows. An internal region of the *pyrF* gene of *P. putida* (*PP_1815*, orotidine 5’- phosphate decarboxylase) was cloned into a vector that cannot replicate in Pseudomonas species (**Fig. 3a**). Integration of the entire pTP·Δ*pyrF* plasmid in the target locus of strain EM173 blocked the PyrF activity altogether. Since this enzyme catalyzes the last essential step in the de novo biosynthesis of pyrimidines (53), the resulting *P. putida* insertion mutant is rendered auxotroph for uracil (Ura). The extent of DNA recombination brought about by selected factors can be assessed in this strain by scoring the reversion to prototrophy due to excision of the plasmid DNA inserted in the *pyrF* locus. The 2,4-DNT degradation pathway was integrated in the chromosome of this reporter strain as previously indicated (54), giving rise to *P. putida* EM·DNT·U (i.e., EM173 *dntABDEG dntR* Δ*pyrF*, Ura–). *P. putida* EM·DNT·U was challenged with 2,4-DNT at 0.5 mM, and cells were plated in M9 minimal medium with or without Ura. The frequency of appearance of *P. putida* prototroph (Ura+) clones was quantified (**Fig. 3b**), and the strain degrading 2,4-DNT showed a 4-fold increase in the frequency of recombination as compared to control conditions. Norfloxacin (NOR) was used as a positive control. NOR is a fluoroquinolone that interferes with the maintenance of chromosomal topology by targeting DNA gyrase and topoisomerase IV, trapping these enzymes at the DNA cleavage stage and thereby preventing strand rejoining (55). Introduction of double-stranded DNA breaks following topoisomerase inhibition by NOR thus induces the SOS response (37, 56). In the presence of NOR, we detected a 7-fold increase in DNA recombination using our reporter system (**Fig. 3b**).

**Fig. 3.**
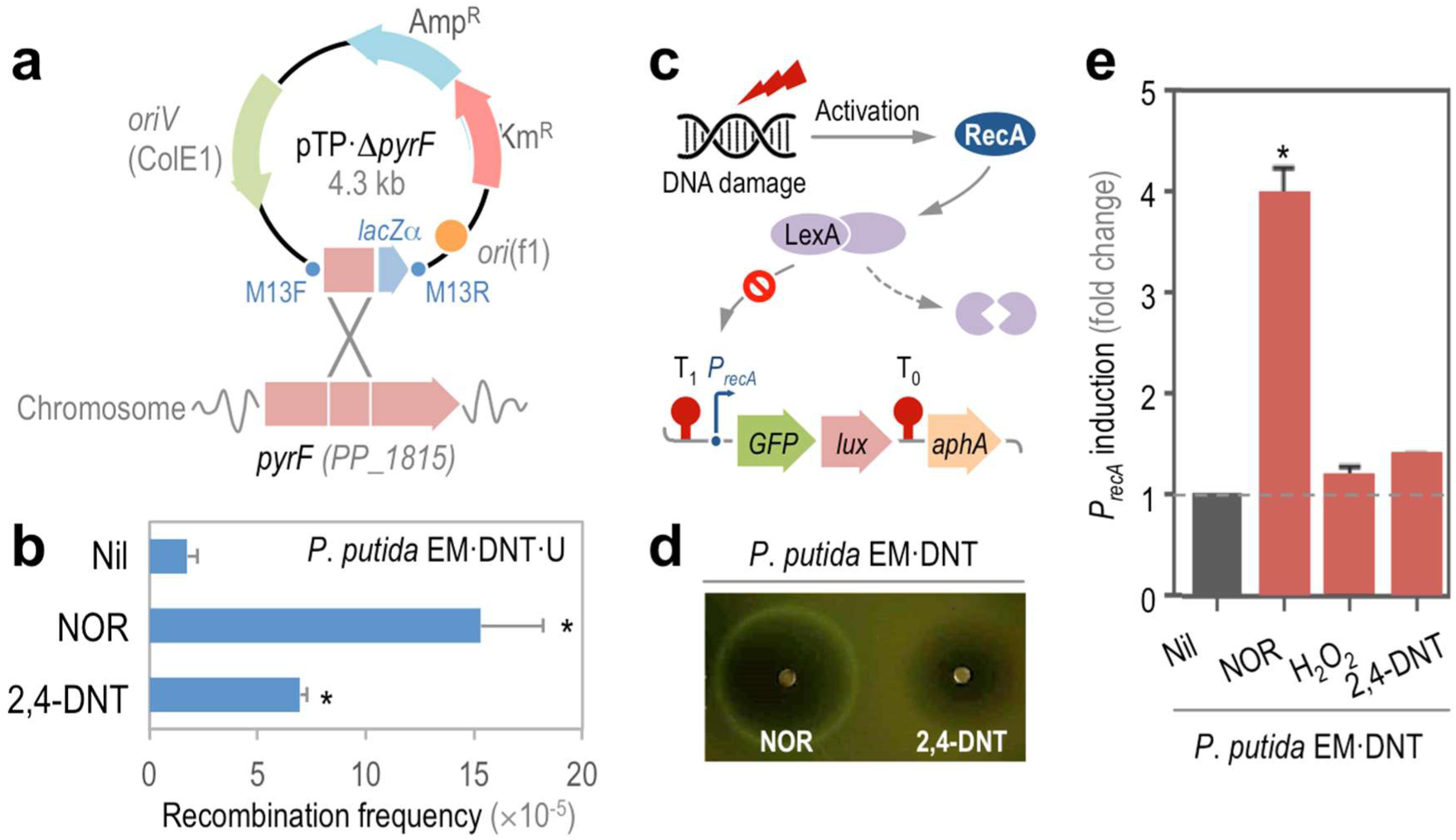
Effects of 2,4-DNT degradation on DNA recombination and SOS response. **(a)** Construction of a reporter *P. putida* strain to study DNA recombination. A suicide, integrative plasmid, obtained as detailed in the Supplementary Information, was used to disrupt the *pyrF* gene of *P. putida*, which results in uracil auxotrophy. The frequency of excision of the plasmid from the chromosome of *P. putida* EM·DNT·U (i.e. reversion to prototrophy) was adopted as an indication of DNA recombination. **(b)** DNA recombination frequency upon exposure to 2,4-DNT or norfloxacin (NOR). Bars represent mean values ± SD (n = 4), and the asterisk identifies significant differences at *P* < 0.05 as indicated by the Student’s t test. **(c)** General mechanism of SOS response upon DNA damage. The RecA protein, stimulated by either damaged or single-stranded DNA, triggers the inactivation of the LexA (a repressor of the SOS response genes) thereby inducing the response. The LexA degradation-dependent activation of the recA promoter of *P. putida* was used as a proxy of the SOS response by constructing an msf·GFP-based biosensor. Elements in the outline are not drawn to scale. **(d)** Testing of the SOS response-biosensor in soft-agar experiments. *P. putida* EM·DNT was transformed with the reporter plasmid, and two filter paper discs, soaked with either NOR or 2,4-DNT, were applied onto the bacterial lawn. The plates were photographed under blue light after 24 h of incubation. **(e)** Quantification of the SOS response-biosensor activity in cultures of *P. putida* EM·DNT in the presence of the additives indicated. Bars represent the mean value of the fold-change in msf·GFP fluorescence ± SD (*n* = 4), and the asterisk identifies significant differences at *P* < 0.05 as indicated by the Student’s t test. The baseline in cultures with no additives (Nil) is indicated with a dashed gray line.

Since degradation of 2,4-DNT brings about an increase in the frequency of DNA recombination, we wondered whether this would result in the activation of the SOS response (i.e. DNA damage response), that is stimulated by single- or double-stranded breaks in genomic DNA (57). When DNA is injured, the RecA protein binds to DNA in the damaged region to form a filament. This filament interacts with a dimer of the LexA transcriptional repressor, activating its self-cleavage and causing the dissociation of LexA from its targets and inducing the SOS regulon (**Fig. 3c**). The promoter of *recA* is one of the targets of LexA (58), an occurrence that was exploited in this work by constructing a biosensor of RecA activity. The promoter region of recA (*PP_1629*), including the putative LexA binding site, was cloned in front of a promoter-less *GFP-luxCDABE* dual reporter system (59). The resulting reporter plasmid was introduced into *P. putida* EM·DNT, and the system was tested by exposing the cells to NOR and 2,4- DNT in a qualitative soft-agar diffusion test (**Fig. 3d**). Exposure of strain EM·DNT carrying the RecA reporter to NOR resulted in a distinct halo of growth inhibition, the boundaries of which gave off a strong GFP signal when observed under UV light. 2,4-DNT, in contrast, did not seem to elicit a similar response in the cells carrying the RecA reporter. A similar pattern was observed in the GFP-dependent fluorescence when the experiment was repeated in liquid cultures of *P. putida* EM·DNT bearing the SOS response biosensor, as quantified by flow cytometry (**Fig. 3d**). Addition to NOR at sub-inhibitory concentrations (250 ng ml-1) caused a 4-fold induction of the RecA reporter, whereas neither 2,4-DNT nor H2O2 resulted in any significant activation of the SOS response. Taken together, these results indicate that degradation of the aromatic substrate promotes DNA recombination extent, without significantly affecting the activity of the SOS response. The next question was whether other forms of mutagenesis, such as point mutations in genomic DNA, could be also correlated to 2,4-DNT biodegradation by the engineered *P. putida* strain.

### Biodegradation of 2,4-DNT in *P. putida* barely increases DNA mutagenesis

Although ROS production stemming from 2,4-DNT metabolism in *P. putida* EM·DNT does not trigger significantly a SOS response and only stimulates recombination to a moderate degree, it was still possible that oxidative damage to DNA, generation of 8-oxoG and general stress-induced relaxation of mismatch repair could increase the overall level of DNA mutagenesis. To inspect this possibility (which was observed in *Burkholderia* sp. R34) the mutagenesis rate was assessed by exposing *P. putida* EM·DNT cells to 2,4-DNT and counting the number of rifampicin rifampicin-resistant (Rif^R^) colonies after plating on a solid culture medium. The antibacterial effects of Rif are based on its ability to bind the β subunit of the RNA polymerase, thereby blocking the elongation of the nascent RNA molecule. Rif^R^ clones usually harbor RpoB mutations in amino acid residues that make contact with Rif (60), rendering RNA polymerase insensitive to the antibiotic (61, 62). Under our experimental conditions, the background rate of RifR clones (i.e., incubating *P. putida* EM·DNT cells in the presence of DMSO) was *M* = (9.1 ± 0.7)×10^-9^. Exposure of *P. putida* EM·DNT cells to 2,4-DNT increased this level of mutagenesis by a mere 28% (and the difference with control experiments was not statistically significant). This was unexpected, as high levels of endogenous ROS are translated onto a mutagenic state in other bacteria. That this was not the case in the strain constructed in this study indicated that the molecular mechanisms that connect ROS-stress with DNA mutagenesis in *P. putida* are plausibly different from those operating in other bacterial species and other factors could be at play. Yet, what could be such factors? One metabolic signature that characterizes *P. putida* is its highly reductive redox metabolism. We thus wondered next whether the interplay between ROS production and DNA mutagenesis could be shielded by such a background metabolism that is so characteristic of this species.

### Altering the redox status of *P. putida* increases the mutagenic effect of the 2,4-DNT degradation pathway

*P. putida* maintains an adequate supply of reducing power through the activity of the EDEMP cycle (44), a core metabolic architecture that combines individual biochemical steps from the Entner-Doudoroff, the Embden-Meyerhof-Parnas, and the pentose phosphate pathways (**Fig. 4a**). By recycling part of the pool of trioses-phosphate back to hexoses-phosphate, this metabolic cycle mediates NAD(P)H formation and enables catabolic overproduction of reduced pyridine nucleotides. This evolutionary-driven metabolic occurrence helps explaining the very high resistance to environmental insults (e.g. oxidative stress) displayed by *P. putida* (63-65). Moreover, antioxidant responses in bacteria exposed to xenobiotics rely on the generation of reducing power that the cells use to counteract ROS formation. For instance, AhpC, a hydroperoxide detoxifying enzyme (**Fig. 2c**), is reduced by peroxiredoxin reductase (AhpF), a process that requires NADH (66). On this basis, and taking into consideration that [i] 2,4-DNT degradation results in the generation of ROS and [ii] the oxidative damage caused by ROS stimulates DNA damage, we set to explore the relationship between redox status and DNA mutagenesis in *P. putida* EM·DNT. To investigate this issue, the redox status of *P. putida* EM·DNT was artificially perturbed by altering the intracellular availability of reduced nicotinamide cofactors through conditional expression of the nox gene of Streptococcus pneumoniae (67, 68). Nox is a NADH-specific oxidase enzyme from that converts O_2_ to H_2_O with a negligible formation of H_2_O_2_ (**Fig. 4b**). Cellular energy demands are not affected by Nox, and thus nox expression allows for the specific investigation of the impact of NADH oxidation without affecting the overall fitness of the cells (69). A synthetic *NADH burning* device was designed for this purpose, in which the gene encoding NADH oxidase from S. pneumoniae was placed under control of the orthogonal ChnR/P_chnB_ expression system (**Fig. 4c**), inducible upon addition of cyclohexanone (70, 71). We tested the effect of NADH oxidase on the overall physiology of *P. putida* EM·DNT by measuring the specific Nox activity in cell-free extracts and evaluating growth rates on succinate cultures (**Fig. 4d**). Induction of the synthetic device by addition of 0.1 mM cyclohexanone resulted in a specific Nox activity of 3.9 ± 0.6 μmol ·mg_protein_^−1^ · min^−1^, ca. 40-fold higher than the background oxidase activity in the cell-free extract of the control strain transformed with the empty vector. Under these conditions, the specific growth rate of *P. putida* EM·DNT was reduced by ca. 30%. As a direct indication of the role of Nox in mediating a redox imbalance, the reduced nucleotide content was evaluated in cell-free extracts of *P. putida* EM·DNT. The NADH concentration was 176 ± 25 and 103 ± 38 μM for the strain carrying the empty pSEVA2311 vector or plasmid pS2311·Nox, respectively, demonstrating that the overexpression of *nox* in *P. putida* cells decreases the reducing power content. The next step was to evaluate if an alteration in the NADH content affects DNA mutagenesis.

**Fig. 4.**
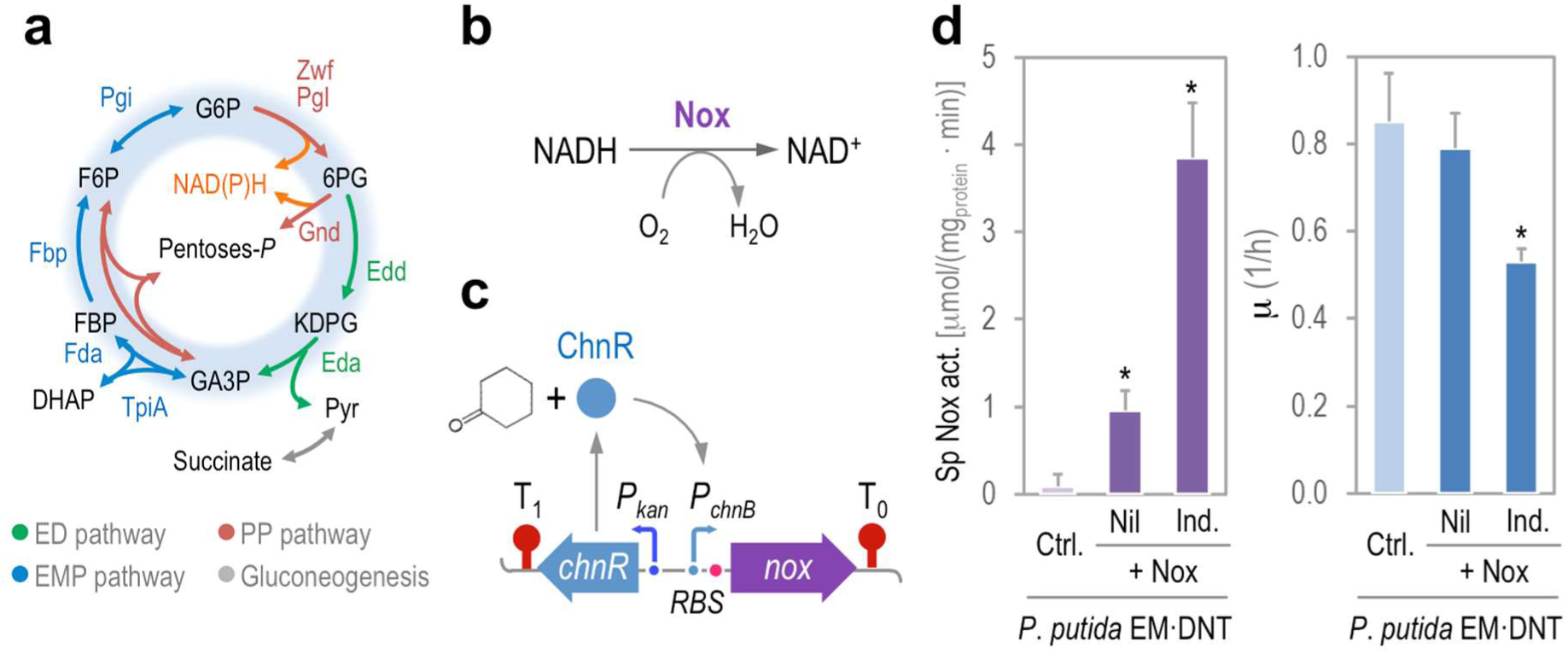
Perturbation of the redox metabolism of *P. putida*. **(a)** Simplified scheme of the upper carbon metabolism of *P. putida*. Note that redox balance is maintained through the action of the EDEMP cycle, indicated in blue shading in the diagram. Abbreviations: ED pathway, Entner–Doudoroff pathway; EMP pathway: Embden–Meyerhof–Parnas pathway; PP pathway, pentose phosphate pathway; G6P, glucose-6-*P*; F6P, fructose-6-*P*; FBP, fructose-1,6-*P_2_*; DHAP, dihydroxyacetone-*P*; GA3P, glyceraldehyde-3-*P*; 6PG, 6-phosphogluconate; and KDPG, 2-keto-3-deoxy-6-phosphogluconate. **(b)** Reaction catalyzed by the water-forming Nox NADH oxidase from Streptococcus pneumoniae. **(c)** Construction of a synthetic NADH burning device for tightly-regulated expression of *nox*. The gene encoding Nox was placed under the control of the PchnB promoter, which responds to the cyclohexanone-activated ChnR regulator. Elements in the outline are not drawn to scale. **(d)** Nox activity and impact of endogenous redox imbalance in the overall physiology of *P. putida* EM·DNT. The specific (Sp) *in vitro* Nox activity was compared in *P. putida* EM·DNT carrying either the empty pSEVA2311 vector (Ctrl.) or plasmid pS2311·*Nox*, with (Ind.) or without (Nil) addition of cyclohexanone at 0.1 mM to induce the expression of nox. The specific growth rate (μ) was determined in the same cultures. Each bar represents the mean value of the corresponding parameter ± SD (*n* = 5), and the asterisk identifies significant differences at *P* < 0.05 as indicated by Student’s *t* test.

*P. putida* EM·DNT was transformed with the nox-expressing plasmid or the empty pSEVA2311 vector, and the resulting strains were exposed to different combinations of 2,4-DNT and cyclohexanone. The rate of mutagenesis *M* was explored by assessing the appearance of Rif^R^ clones after each treatment (**Fig. 5a**). Exposure of the cells to 2,4-DNT or activation of Nox alone did not result in a significant increase in DNA mutagenesis. The combination of the two treatments, however, produced a 6-fold increase in *M*, indicating that the redox imbalance introduced by Nox exacerbates the mutagenic impact of 2,4-DNT. Yet, what is the nature of the mutations that lead to RifR in such redox-stressed scenario?

**Fig. 5.**
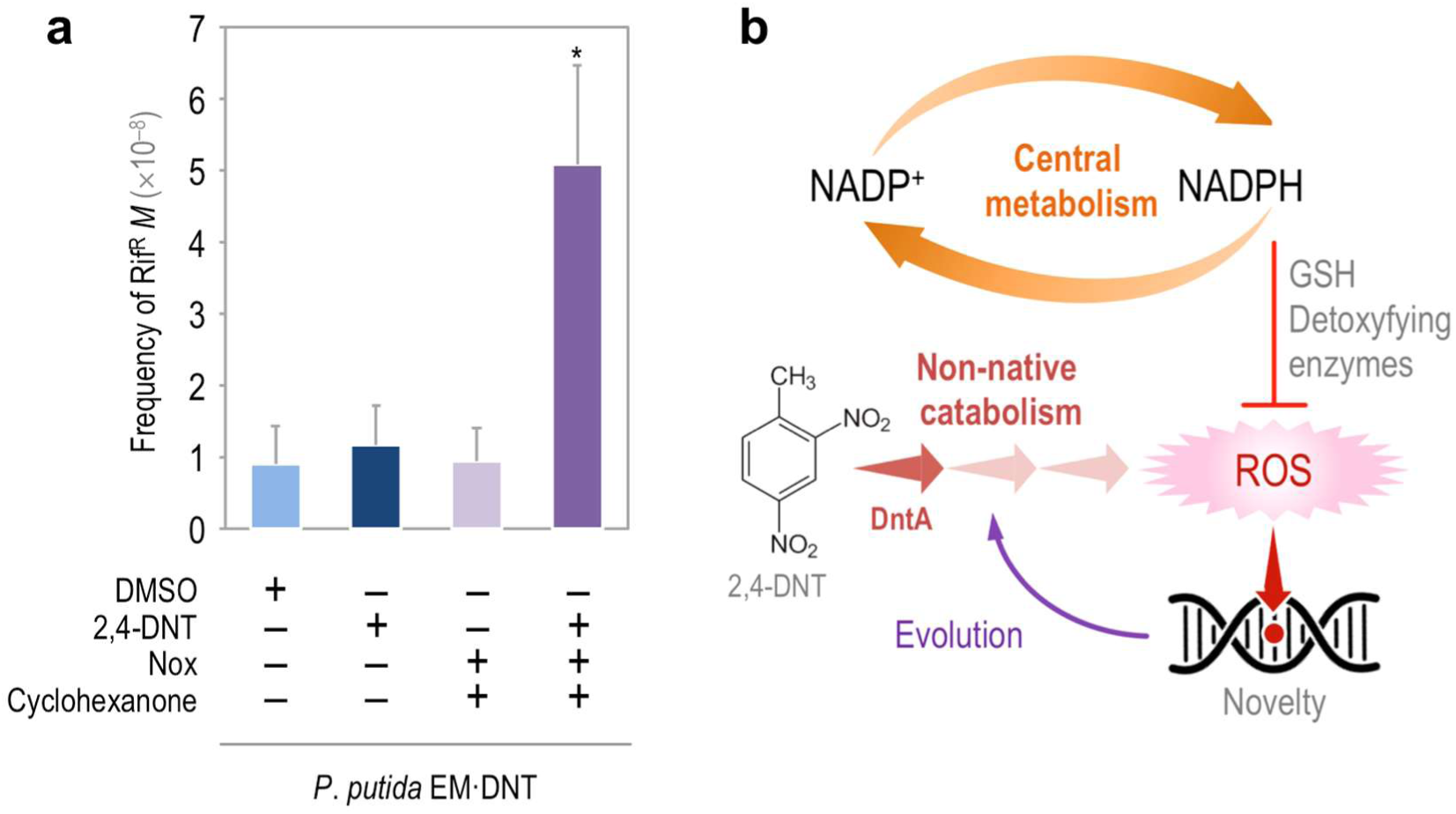
DNA mutagenesis in redox-challenged *P. putida* EM·DNT facing 2,4-DNT. **(a)** Frequency of spontaneous rifampicin-resistant mutants (*M*) in cells exposed to 2,4-DNT at 0.5 mM, DMSO (the 2,4-DNT solvent carrier), cyclohexanone at 0.1 mM (inducer of the synthetic device driving the expression of the Nox NADH oxidase), and combinations thereof. The dimensionless frequencies of mutation were calculated using the Ma-Sandri-Sarkar maximum likelihood estimator. Bars represent the mean M values ± SD (*n* = 4), and the asterisk identifies significant differences at P < 0.05 as indicated by the Student’s *t* test. **(b)** Model for metabolism-driven evolution. Faulty redox reactions on novel substrates originate ROS, which may cause direct or indirect damage to DNA in a fashion dependent on the background metabolism, which provides the reducing power necessary to fuel detoxifying enzymes (typically dependent on reduced glutathione, GSH).

### The spectrum of *rpoB* mutations depends on the intracellular redox status

Mutations in *rpoB* that confer Rif^R^ are well conserved among prokaryotes, and the amino acid changes that produce this phenotype can be grouped in three clusters in a central region of RpoB (61). We sequenced ca. 100 clones per experimental condition to obtain the spectrum of mutations in each case (**Table 1**). All the mutations detected in this analysis were located in Cluster I of RpoB [between nucleotides 1,528 and 1,626 of *rpoB*, as defined by Jatsenko et al. (61)], and they could be grouped into eight categories. Five of the categories were transitions (i.e., A→G and C→T), two of them were transversions (i.e., A→T and C→A), and the rest of the *rpoB* mutations found in Rif^R^ clones were grouped as *Other* (a category that includes the less abundant *rpoB* mutations). The two predominant mutations found resulted in Q518L and D521G changes in RpoB, and their frequency of appearance did not suffer any significant change across the experimental treatments tested herein. Interestingly, the frequency of C→A transversion that generates a L538I change in RpoB was influenced by both exposure to 2,4-DNT (2- fold increase) and the activation of Nox (4-fold increase). The effect of the two treatments was additive, further linking biodegradation of the substrate and redox status of the cells with DNA mutagenesis. C→A transversions are known to be caused by the presence of 8-oxoG in the DNA (72), which results from the attack of ROS to purine residues. This result thus indicated that oxidative stress caused by 2,4-DNT degradation ends up in oxidation of purines (as observed in *Burkholderia* sp. R34), but also shows that the more reductive redox status of *P. putida* provides effective protection against this mutagenic effect. Other mutations arising by exposing the cells to 2,4-DNT have a less clear origin, e.g. the activity of the widely distributed Y-family of DNA polymerases (73). Members of this family are characterized not only by their ability to replicate damaged DNA, but also by their lack of processivity and low fidelity when copying undamaged template. In *P. putida* KT2440, this protein category is represented by DinB (PP_1203) and the error-prone DNA polymerase DnaEB (PP_3119) (74). In sum, the spectrum of *rpoB* mutations detected in our study suggests that ROS-induced mutagenesis triggered by 2,4-DNT metabolism merges direct damage to DNA caused by misincorporation of 8-oxoG with other stress-response mechanisms that relax fidelity of DNA replication and/or prevent repair.

**Table 1.** Nature and frequency of mutations in *rpoB* (*PP_0447*) conferring resistance to rifampicin^*a*^.

| Mutation | *P. putida* EM-DNT carrying |  |  |  |  |  |
| --- | --- | --- | --- | --- | --- | --- |
|  | pSEVA2311 (empty vector) |  |  | pS2311-Nox (NADH oxidase) |  |  |
|  | DMSO | 2,4-DNT | 2,4-DNT | DMSO | 2,4-DNT | 2,4-DNT |
|  | (control) |  | +<br>cyclohexanone | (control) |  | +<br>cyclohexanone |
| Q518L (A→T) | 31 | 27 | 29 | 32 | 19 | 15 |
| Q518R (A→G) | 14 | 11 | 13 | 11 | 11 | 10 |
| D521G (A→G) | 21 | 25 | 22 | 19 | 26 | 24 |
| H531R (A→G) | 12 | 7 | 5 | 15 | 12 | 6 |
| H531Y (C→T) | 8 | 9 | 10 | 7 | 4 | 8 |
| S536F (C→T) | 9 | 4 | 3 | 10 | 8 | 9 |
| L538I (C→A) | 1 | 11 | 13 | 5 | 11 | 21 |
| Other | 4 | 6 | 5 | 1 | 9 | 7 |

<sup>a</sup> The percentage of amino acid changes in RpoB is indicated along the DNA mutation that originated the substitution. These percentages were calculated by sequencing an internal region of *rpoB* in at least 90 rifampicin-resistant clones for each experimental condition. The mutations indicated in the list were selected since they were observed across all the experimental conditions tested. The remaining point mutations are grouped as "Other". Abbreviations: DMSO, dimethyl sulfoxide; and 2,4-DNT, 2,4-dinitrotoluene.

## DISCUSSION

Evolvability of bacteria for conquering novel nutritional and physicochemical landscapes is the result of the interplay between endogenous and exogenous factors. Propagation of new pathways through a bacterial population is fostered by different mechanisms of horizontal gene transfer, e.g. conjugation, transformation, and phage- or ICE^1^-mediated DNA acquisition (75, 76). In this case, the emergence of genetic and functional diversity largely relies on the balance between the effectiveness of the transfer events and the ability of the recipient bacteria to gain an advantage thereof. Not surprisingly, microbial species with a high capacity to receive foreign DNA usually evolve new traits (77)—sometimes as undesirable as antibiotic resistances. However, horizontal gene transfer does not account for the onset of fresh routes for xenobiotic catabolism, which require mutation of pre-existing pathways to reshape their substrate range and rewire their biochemical outputs to the existing metabolic network. Since these changes must necessarily occur in single hosts, one can safely predict that the higher mutagenic regime they have, the faster the solution space will be explored, and an optimum eventually found. Two common triggers of such mutagenic regime include carbon starvation and endogenous production of ROS (78-81). Given that these mechanisms are virtually universal through all aerobic bacteria, the question arises on why most of aerobic routes for degradation of xenobiotic aromatics involve the action of Rieske non-heme iron oxygenases borne by Pseudomonads and closely related bacteria. The results described in this article provide some rationale for this longstanding observation. The mechanism of this type of oxygenases at their reaction center involves activation of an oxygen molecule by a Fe-S cluster to make the oxygen reactive towards the aromatic substrate (82, 83). Even with a good enzyme-substrate fit, purely stochastic effects at the atomic level may result in coupling failure and release of ROS, the discharge of which will increase if the fitting of the substrate to the enzyme is worse (24-27). We have argued before that, because of its mutagenic potential, the release of ROS is a major driver of the evolution of the cognate bacterial population towards a new biodegradative target (34). Yet, why are such biodegradative abilities so conspicuous in Pseudomonads? The data above suggest that, while ROS production during 2,4-DNT degradation is unavoidable, DNA mutagenesis is ultimately controlled by the endogenous redox status of the corresponding cells. The highly reductive NAD(P)H regime associated to the distinct central carbon metabolism of *P. putida* thus seem to render this species well suited for hosting pathways involving strong redox transactions (**Fig. 5b** and **6a**). At the same time, a dearth in NAD(P)H availability would result in a high DNA mutagenesis regime. Although data supporting this notion have been generated in only two environmental bacteria (**Pseudomonas putida** and *Burkholderia* sp.), it is likely that they are specific cases of a general evolutionary principle that is reminiscent of the concept of *anti-fragility* (34, 84) or *hormesis*, i.e. beneficial effects of a low-dose of a toxic input (85, 86), observed in other biological systems. Under this frame, there seems to be an optimum in the diversity-generating regime caused by stress (e.g. ROS) that is enough to generate sufficient evolutionary bet-hedging for coping with a new nutrient of physicochemical condition while keeping the thereby diversified—but also damaged population—within viability (**Fig. 6b**). Is within this solution space that bacterial cells equipped with novel catabolic pathways for alien substrates are to be found. The data above suggest that such a window of optimality is shaped by the background metabolic network, specifically by the redox regime. ROS formation has been observed in other environmental bacteria running biodegradation pathways for xenobiotics (87-90), and it is plausible that similar evolutionary processes apply in other instances.

**Fig. 6.**
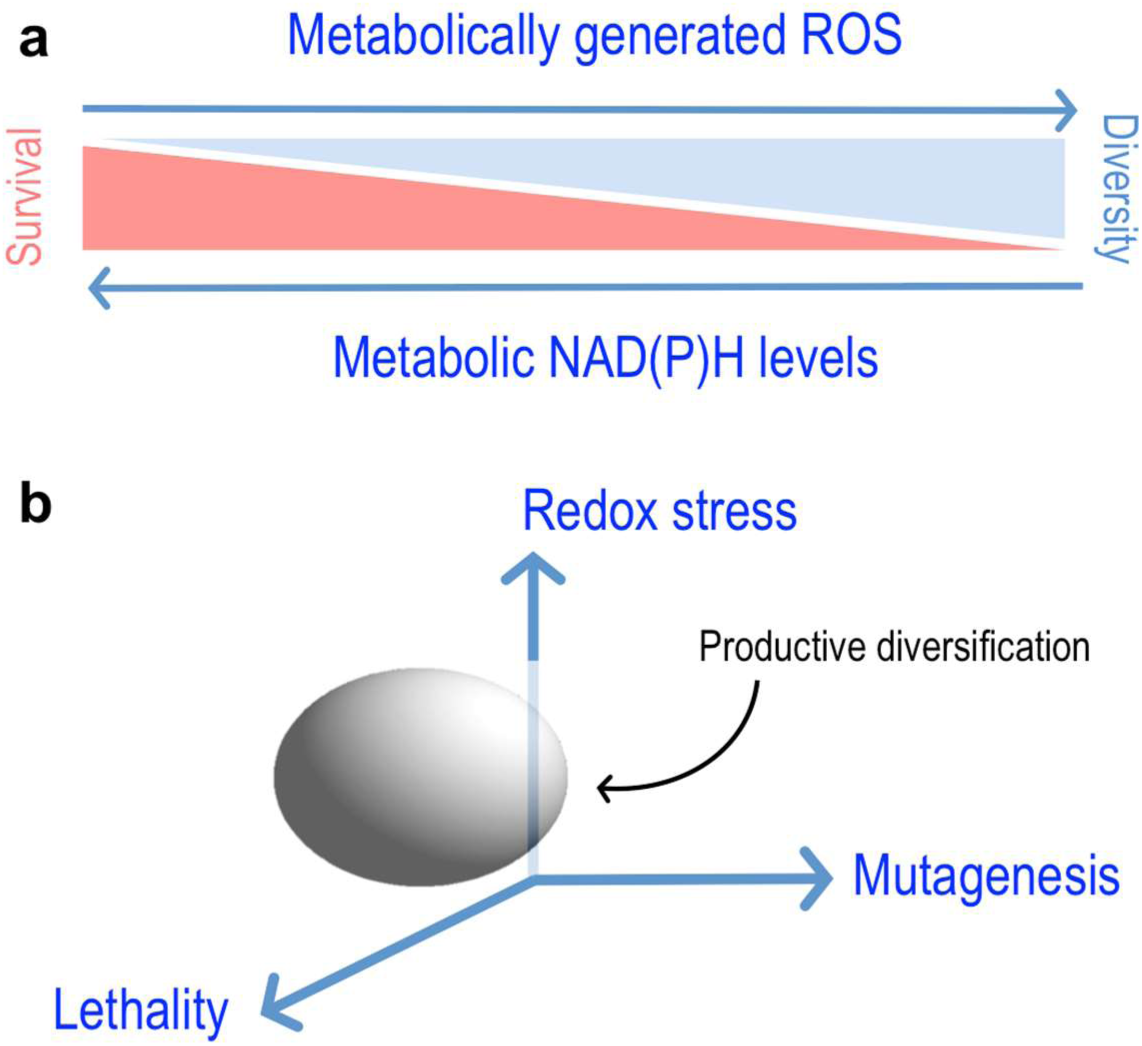
Evolutionary hormesis driven by metabolic production of ROS. **(a)** Interplay between redox stress and genetic and functional diversification of bacteria. As explained in the text, the action of ROS as a direct or indirect DNA mutagenesis agent is checked by the endogenous levels of reductive currency [i.e. availability of NAD(P)H]. This situation defines a scenario in which the gain of diversity occurs at the expense of population survival. **(b)** Species-specific definition of a productive diversification space. In this work, we argue that the *evolvability* of given bacterial species and the potential to host novel catabolic pathways is framed by the ability of bacteria to endure ROS-driven stress in a fashion that depends on their particular redox metabolism. These independent elements demarcate a space of productive diversification, i.e. an optimum in the *novelty supply* rate of the system (100), that fosters the exploration of the evolutionary solution space which is unique for each bacterial species.

That the redox metabolic status (either inherent or transient) controls evolvability of selectable traits in bacteria raises some corollaries worth to follow up. For instance, ROS may accelerate the emergence of new antibiotic resistances in species subject to endogenous oxidative stress but endowed with a weaker redox metabolism. This opens the possibility of targeting redox biochemistry of virulent strains with drugs that perturb the NAD(P)^+^/NAD(P)H balance. Moreover, our data suggest that some members of a given microbial community could be more innovative than others on the basis of their inherent redox metabolism. There could be innovation *specialists* that nurture genetic diversification and pass the results to less mutable members of a multi-species bacterial consortium through horizontal gene transfer. Finally, ROS-mediated genetic innovation could be repurposed for the sake of generating new pathways through directed evolution or for the optimization of biocatalytic routes for metabolic engineering. In any case, the connection between metabolism and evolvability should not be ignored when addressing the emergence of new traits whether for fundamental understanding of evolutionary processes or for designing new whole-cell biocatalysts.

## MATERIALS AND METHODS

### Bacterial strains and culture conditions

Bacterial strains are listed in Table S1 in the Supplemental Material. *Burkholderia* sp. R34 is a 2,4-DNT degrading specimen (32). *P. putida* EM173 was used as the host of the *dnt* pathway (48). E. coli strains DH5α, CC118, CC118λ*pir*, and HB101 were employed for DNA cloning procedures. Bacteria were grown in rich LB medium or in M9 minimal medium with sodium succinate added at 0.4% (wt/vol) with rotary agitation at 170 rpm (91). *P. putida* was grown at 30°C while E. coli was grown at 37°C. When required, 2,4-DNT was added at 0.25 or 0.5 mM from a 0.5 M stock solution freshly prepared in DMSO. Control experiments were added with an equivalent volume of DMSO. Overnight-grown *P. putida* in M9 minimal medium and succinate was used as inoculum by diluting the bacterial suspension to a starting OD_600_ of 0.05. Cells were grown in the same culture medium until OD_600_ of 0.5, at which point the cell suspension was split: one culture served as a control, and the other one was challenged with 2,4-DNT. Other additives were as indicated in the text, and samples were periodically taken as specified. Additives were added at the following final concentrations: ampicillin, 150 μg ml^−1^ for E. coli or 500μg ml^−1^ for **P. putida*;* chloramphenicol, 30 μg ml^−1^; kanamycin, 50 μg ml^−1^, streptomycin, 80 μg ml^−1^; gentamicin, 10 μg ml^−1^; uracil, 20 μg ml^−1^; H_2_O_2_, 1.5 mM; and cyclohexanone, 0.1 mM. Norfloxacin was used at 25 ng ml^−1^ for stimulation of DNA recombination and at 250 ng ml^−1^ for reporter experiments. All solid media contained 15g l^−1^ agar.

### General DNA techniques and construction of recombinant *P. putida* strains

DNA manipulations followed routine laboratory techniques (91). Details on the design and construction of bacterial strains and plasmids can be found in the Supplemental Material. Plasmids were introduced into *P. putida* strains by electroporation (92, 93).

### Determination of mutation frequencies and detection of rpoB mutations

*P. putida* cultures were grown overnight with the appropriate antibiotics, diluted 1:10,000 into 50 ml of fresh culture medium (initial OD_600_ of 0.05) in 250-ml Erlenmeyer flasks, and incubated for 4 h. These cultures were diluted 1:4 into fresh medium (1 ml in 10-ml test tubes), and incubated for 16 h in the presence of relevant additives. Appropriate dilutions were spread on LB medium plates containing 50 μg ml^−1^ Rif. Colonies were counted after 48 h at 30°C. Luria-Delbrück fluctuation analysis was performed to study the impact of additives in the mutation rate M. M values were computed using the Ma-Sandri-Sarkar maximum likelihood estimator (ratio between the number of Rif^R^ colonies and the total viable count, corrected by plating efficiency) from at least six independent experiments per condition (94, 95). The statistical significance of mean M values across experiments was evaluated with the Student’s t-test. To identify the nature and frequency of point mutations in *rpoB*, ca. 100 Rif^R^ colonies were randomly collected from at least four independent experiments, and the locus was sequenced with primers indicated in the Supplemental Material. The BioEdit^TM^ software (BitSize Bio) was used for sequence comparison.

### Analytical procedures

The ROS-sensitive green fluorescent dye H_2_DCF-DA (Sigma-Aldrich) was used to quantitate ROS formation. Flow cytometry, preparation of cell-free extracts, metabolite extraction, and enzymatic assays followed protocols from our laboratory (45, 46, 93, 96-99), and further details can be found in the Supplemental Material.

### Statistical analysis

All reported experiments were independently repeated at least three times (as indicated in the figure legends), and mean values of the corresponding parameter and standard deviation (SD) is presented. The significance of differences when comparing results was evaluated by means of the Student’s t test.

## ACKNOWLEDGEMENTS

The authors wish to thank J. Blázquez (Spain), M. H. H. Nørholm (Denmark), A. Smania (Argentina), and L. Eltis (Canada) for inspiring discussions. O.A. was supported by TUBITAK-BIDEP through the International Postdoctoral Research Scholarship Programme 2219. This study was supported by HELIOS Project of the Spanish Ministry of Economy and Competitiveness BIO2015-66960-C3-2-R (MINECO/FEDER), and the ARISYS (ERC-2012-ADG-322797), EmPowerPutida (EUH2020-BIOTEC-2014-2015-6335536), and MADONNA (H2020-FET-OPEN-RIA-2017-1-766975) contracts of the European Union to V.D.L. This study was also supported by The Novo Nordisk Foundation through grant NNF10CC1016517 to P.I.N.

V.D.L., P.I.N., and D.P.P. designed the project and the experimental layout. O.A. performed most of the experiments related to strain construction, oxidative stress, substrate degradation, and mutagenesis.

B.C. constructed strains and plasmids used in this study. P.I.N. and D.P.P. analyzed the data. P.I.N. and V.D.L. wrote the manuscript with further contributions from all authors to data analysis and interpretation of the results. The authors declare that there are no competing interests associated with the contents of this article.

## Footnotes

1 Integrative and conjugative elements

